# Unravelling the genetic basis for the rapid diversification of male genitalia between *Drosophila* species

**DOI:** 10.1101/2020.06.19.161018

**Authors:** Joanna F. D. Hagen, Cláudia C. Mendes, Shamma R. Booth, Javier Figueras Jimenez, Kentaro M. Tanaka, Franziska A. Franke, Luis. Baudouin-Gonzalez, Amber M. Ridgway, Saad Arif, Maria D. S. Nunes, Alistair P. McGregor

## Abstract

In the last 240,000 years, males of the *Drosophila simulans* species clade have evolved striking differences in the morphology of their epandrial posterior lobes and claspers (surstyli). These changes have most likely been driven by sexual selection and mapping studies indicate a highly polygenic and generally additive genetic basis. However, we have limited understanding of the gene regulatory networks that control the development of genital structures and how they evolved to result in this rapid phenotypic diversification. Here, we used new *D. simulans / D. mauritiana* introgression lines on chromosome 3L to generate higher resolution maps of posterior lobe and clasper differences between these species. We then carried out RNA-seq on the developing genitalia of both species to identify the genes expressed during this process and those that are differentially expressed between the two species. This allowed us to test the function of expressed positional candidates during genital development in *D. melanogaster*. We identified several new genes involved in the development and possibly the evolution of these genital structures, including the transcription factors Hairy and Grunge. Furthermore, we discovered that during clasper development Hairy negatively regulates *tartan*, a gene known to contribute to divergence in clasper morphology. Taken together our results provide new insights into the regulation of genital development and how this evolves between species.

## Introduction

To understand the evolution of animal morphology we need to better link genotypic and phenotypic changes. This requires identifying the causative genes, how are they integrated into gene regulatory networks, and how changes in these interactions alter developmental processes and consequently the phenotype (Kittelmann et al., 2018, Nunes et al., 2013, Stern, 2011). There has been great progress in identifying genes that cause changes in animal morphology (reviewed in Martin and Orgogozo, 2013). However, we still lack information on the genes that contribute to changes in quantitative traits, such as organ size, and how they combine to achieve this.

The size and shape of male genital organs evolves rapidly among species, driven by sexual selection (Eberhard, 1985, Eberhard, 2010, Hosken and Stockley, 2004, House et al., 2013, Simmons, 2014). For example, the epandrial posterior lobes and claspers (surstyli) have changed dramatically in size in the *Drosophila simulans* species clade in the last 240,000 years (Garrigan et al., 2012) (Fig. 1A). Both the claspers and posterior lobes play important roles during copulation. The claspers open the female oviscapt through interdigitisation of bristles, and help achieve correct copulatory positioning (Acebes et al., 2003, Jagadeeshan and Singh, 2006, Kamimura and Mitsumoto, 2011, Masly and Kamimura, 2014, Mattei et al., 2015, Robertson, 1988, Yassin and Orgogozo, 2013), while the posterior lobes also contribute to stability during mating by inserting into grooves on the female tergites (Kamimura and Mitsumoto, 2011, Robertson, 1988, Yassin and Orgogozo, 2013).

**Figure 1.**
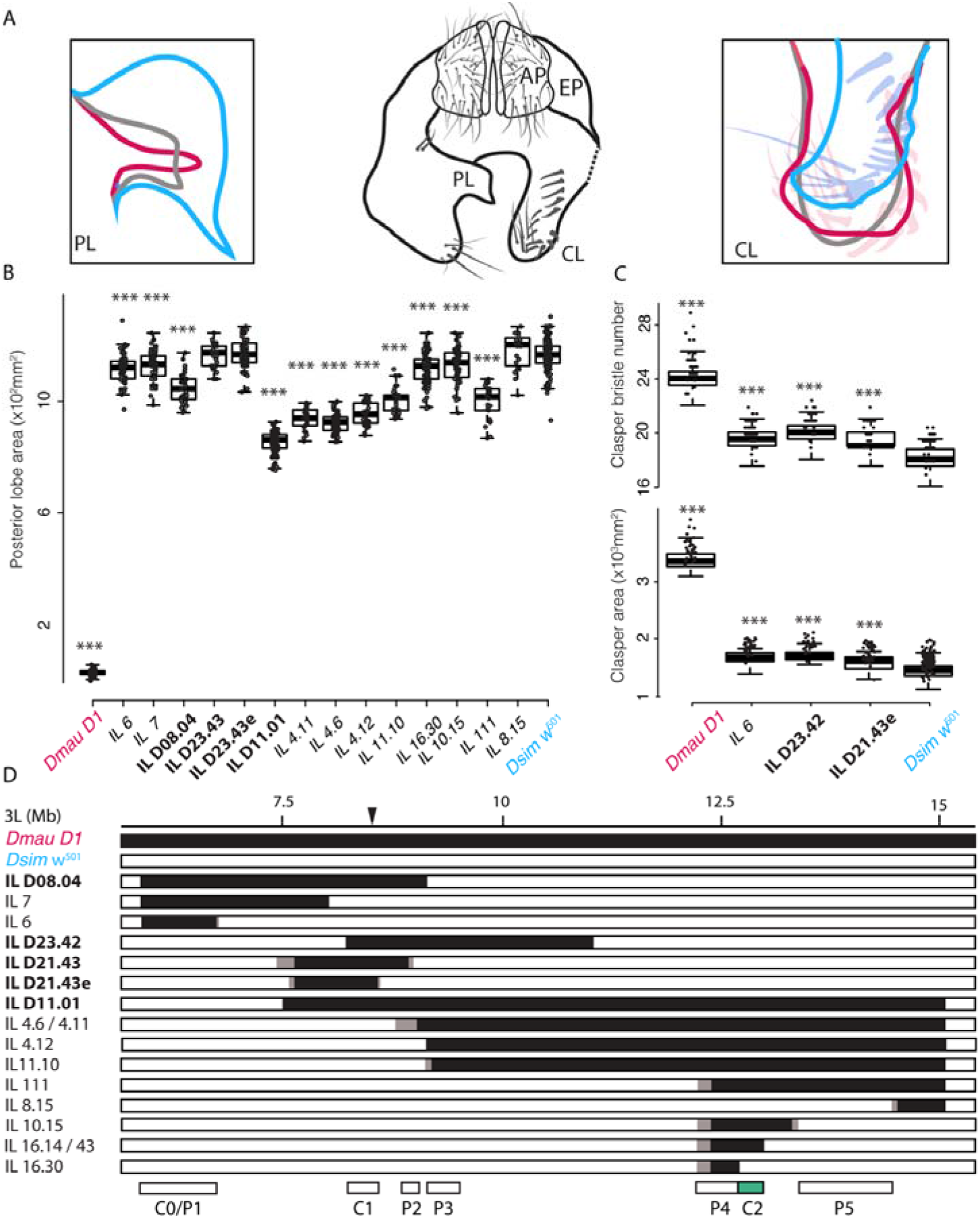
Differences in external male terminal structures among *D. melanogaster* subgroup species and introgression mapping between *D. simulans* and *D. mauritiana*. (A) Schematic representation of a posterior view of *Drosophila* male genital arch. The central diagram depicts the genital arch morphology of *D. melanogaster*. The posterior lobes (left-hand box) typically obscure visualization of the claspers (right-hand box), and therefore they are shown here dissected away on the right-hand-side of the central schematic. The relative size and shape of the lobes and the claspers of *D. melanogaster* (grey), *D. simulans* (blue) and *D. mauritiana* (red) are illustrated in the left and right schematics, respectively. *D. simulans* have smaller claspers, with fewer, thinner and shorter bristles *than D. mauritiana* and *D. melanogaster*. Posterior lobes (PL), claspers (CL), anal plates (AP) and epandrium (EP). (B-C) Mapping and phenotypic effect of candidate regions on posterior lobe size (B), clasper bristle number (C, upper plot) and clasper area (C, lower plot). Boxes indicate the range, upper and lower quartiles and median for each sample. Asterisks indicate significant comparisons with *Dsim w*^501^ where *p* < 0.001 *** (Dunnett’s test for posterior lobe and clasper size, Dunn’s test for clasper bristle number, Supplementary File S2D-S2F). Differences in the effect of the introgressed regions (Supplementary Text) on posterior lobe size (B) and clasper size/bristle number (C), allowed refinement of candidate regions P1 – P5, C0 and C1 (D). The previously identified C2 region is shown in green (Hagen et al., 2019) (D). Black bars indicate *Dmau D1* DNA, white bars indicate *Dsim w*^501^ DNA, and grey boxes regions containing break points that have not been precisely determined. The black triangle indicates the position of P-element insertion originally used for generating the introgressions. New introgression lines are shown in non-bold font.

The posterior lobes are a novelty of the *D. melanogaster* species subgroup (Glassford et al., 2015, Jagadeeshan and Singh, 2006, Kopp and True, 2002). In *D. mauritiana*, they are small, thin finger-like projections in comparison to the much larger, helmet shaped lobes of *D. simulans* (Fig. 1A). *D. melanogaster* has intermediate sized lobes, which are trapezoid shaped (Fig. 1A), while the *D. sechellia* lobes are also intermediate in size and resemble “boots”. It is important to note that there is some variation within species but the extremes of intra-specific variation do not overlap with the differences observed between species (Hackett et al., 2016, McNeil et al., 2011).

The claspers lie underneath the posterior lobes, and are more than twice as large in *D. mauritiana* compared to *D. simulans*, with a third more bristles (Tanaka et al., 2015, True et al., 1997) (Fig. 1A). The morphology of these bristles also differs between the species, with the *D. mauritiana* bristles being generally shorter and thicker than those of *D. simulans* (Tanaka et al., 2015, True et al., 1997). *D. sechellia* male claspers have very similar morphology to those of *D. simulans*, while the claspers of *D. melanogaster* appear to be intermediate between *D. mauritiana*, and *D. simulans*/*D. sechellia* (Fig. 1A).

Genetic mapping of changes to posterior lobe and clasper morphology among *D. melanogaster* subgroup species have shown that these differences are polygenic and generally additive (Coyne et al., 1991, Laurie et al., 1997, Liu et al., 1996, Macdonald and Goldstein, 1999, Tanaka et al., 2015, Tanaka et al., 2018, True et al., 1997, Zeng et al., 2000). For example, up to 19 QTL have been identified for the difference in posterior lobe size between *D. mauritiana* and *D. simulans*, and QTL have been mapped to all major autosomal arms for the differences in clasper size between these species (Laurie et al., 1997, Tanaka et al., 2015, Tanaka et al., 2018, True et al., 1997, Zeng et al., 2000). Therefore, it appears that many loci contribute to these differences in genital organ size.

We previously used an introgression based approach to map QTL on chromosome 3L underlying posterior lobe and clasper size differences between *D. mauritiana* and *D. simulans* (Hagen et al., 2019, Tanaka et al., 2015). The genomes of these lines were *D. simulans*, apart from introgressed regions of *D. mauritiana* DNA on 3L (Hagen et al., 2019, Tanaka et al., 2015). The regions that we found to contribute to posterior lobe and clasper size differences were mutually exclusive; suggesting that different genes underlie divergence in these two structures (Tanaka et al., 2015). Furthermore, this approach revealed that sequence divergence in *tartan* (*trn*), which encodes a leucine-rich repeat transmembrane protein, contributes to the larger claspers of *D. mauritiana* compared to *D. simulans* (Hagen et al., 2019). This is likely due to more widespread and persistent expression of *trn* in the developing claspers in *D. mauritiana* (Hagen et al., 2019). However, since *trn* does not appear to contribute to posterior lobe size differences and explains only 16% of the clasper size difference between the species (Hagen et al., 2019), there must be additional loci involved in posterior lobe and clasper size differences on chromosome 3L.

To try to identify other causative genes on 3L, we generated new introgression lines to further refine existing candidate regions (Tanaka et al., 2015). We complemented this approach with RNA-seq on the developing genitalia of both species to identify genes expressed and differentially expressed both genome-wide and in the mapped regions. Subsequent functional testing of positional and expression candidate genes in *D. melanogaster* identified new candidate genes and novel players involved in genital development, including the TFs *Grunge* (*Gug*) and *hairy* (*h*), which appear to positively and negatively regulate clasper size respectively. Furthermore, we found that H represses *trn* expression in the developing clasper suggesting that changes in this regulatory interaction may contribute to inter-specific differences in this structure. Taken together our findings provide new insights into the genetic interactions that underlie genital development, as well as the divergence of genital morphology between *Drosophila* species.

## Results

### Mapping regions underlying male genital divergence between D. simulans and D. mauritiana

Previously we resolved the C2 candidate region for clasper size divergence between *D. simulans and D. mauritiana* by successfully identifying *trn* as the causative gene in this region (Hagen et al., 2019). In order to increase the resolution of other candidate regions contributing to genitalia divergence (Tanaka et al., 2015), we generated 23 new introgression lines with smaller introgressed *D. mauritiana* regions in a *D. simulans* background (Supplementary File 1). We mapped clasper size and clasper bristle number to two regions that collectively explain 16.8% of clasper size differences between these species (Table 1, Supportive Text). We confirmed the location and effect size of the previously identified C1 (Tanaka et al., 2015) and identified a new region, C0, which explains 11% of the divergence in clasper morphology (Fig. 1D, Table 1, Supportive Text). We mapped posterior lobe size to five regions that collectively explain 29.3% of posterior lobe size differences between *D. mauritiana* and *D. simulans*, two of which, P4 and P5, are new (Fig. 1D, Table 1, Supportive Text). In total, these regions contain 380 proteincoding genes (as annotated in *D. melanogaster*, Table 1).

**Table 1.**
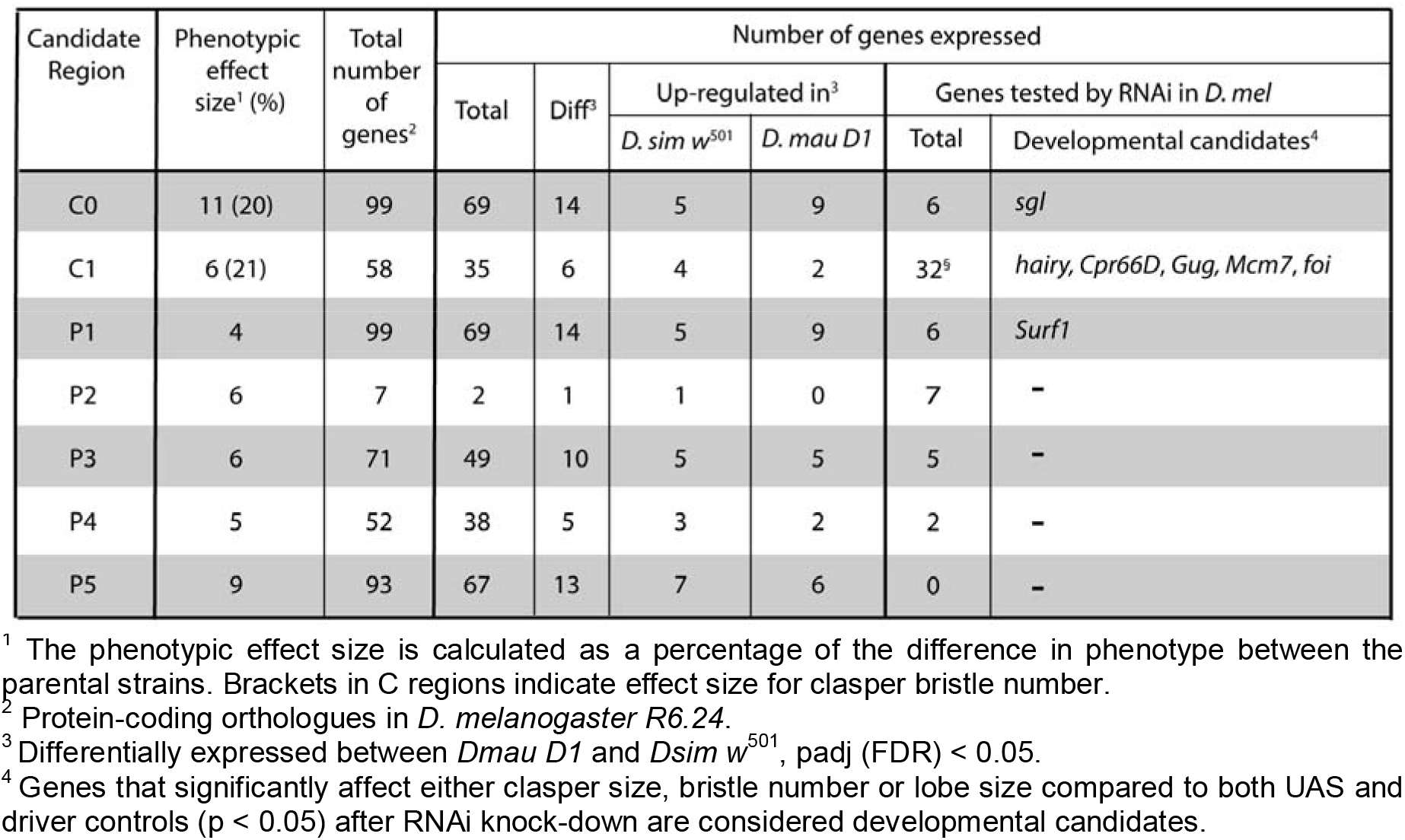
Summary of candidate regions underlying clasper and posterior lobe divergence.

### Analysis of genes expressed in developing male genitalia

We next carried out RNA-seq on developing genitalia of *Dmau D1* and *Dsim w*501. For each species we pooled developing genitalia from stages 2 and 4.5 (for staging see Hagen et al., (2019)). This allowed us to assay the genes expressed in the developing genitalia and those differentially expressed between these two species genome-wide and in our mapped regions.

We detected expression of 8,984 and 8,458 genes above the threshold value of 1 TPM in all biological replicates in the developing genital arches of *Dsim w*^501^ and *Dmau D1*, respectively (Supplementary File 4A). A total of 760 genes are only expressed in *Dsim* w^501^, while 264 genes are only expressed in *Dmau D1*. However, many of these genes (114 and 121 genes, respectively) have low expression in the species where they are detected (< 2 TPM on average between replicates) and therefore are less likely to underlie functional expression differences between species. Gene ontology analysis (GO) of the remaining 676 detected genes uniquely detected in *Dsim w*^501^ indicate the most significant enrichment in genes involved in heme binding (Supplementary File 4B) such as *Cyp4d14, Cyp9b2, Cyp6d5, Cyp6t1, Cyp4g1, Cyp12a5, Cyt-c-d, Cyp6d2, glob2, Cyp6a20* and *Cyp4aa1*. The remaining 143 detected genes exclusive to *Dmau D1* were enriched for ion transmembrane transporters (Supplementary File 4B), the majority of which were ionotropic receptors (IR’s) (*IR76b, IR7g, IR60b, IR7f, IR25a*) and the ionotropic glutamate receptor *Eye-enriched kainate receptor* (*Ekar*).

Of the 8194 genes detected in both species, 1169 were significantly differentially expressed between *Dsim w*^501^ and *Dmau D1*, with 547 upregulated in the former and 622 in the latter respectively (padj < 0.05, Supplementary File 4A).

To explore divergence in gene regulation in the developing male genitalia of these species further we next assessed the expression of transcription factor (TF) encoding genes. We found 802 out of 994 genes encoding TFs and co-factors are expressed in the developing genitalia of *Dmau D1* and *Dsim w*^501^ according to our RNA-seq dataset (Supplementary File 4C). We identified eight TF genes that appear to be exclusively expressed in the developing male genitalia of *Dmau D1*, while sixteen appear to be exclusive to *Dsim* w^501^. However, three and ten of these TF, respectively, were detected at relatively low levels (TPM < 2) and are therefore not likely to contribute to functional regulatory differences in genital development between species. Of the 778 TF genes expressed in both species (Supplementary File 4C), 49 are differentially expressed with 33 upregulated in *Dmau D1*, and sixteen upregulated in *Dsim w*^501^ (Supplementary File 4D).

We then focussed on which of the genes in our mapped introgression regions are expressed in the developing genitalia. We found that 260 of the 380 protein-coding genes located in the introgression mapped regions could be detected in our RNA-seq data, including 31 TFs (Supplementary File 5). Fifty of the expressed candidate genes are differentially expressed between *Dsim w*^501^ and *Dmau D1*, with exactly half the genes being up-regulated in each species (Table 1). This includes one TF that is upregulated in *Dsim w*^501^ (*mirror* (P4)) and four in *Dmau D1 (Meiotic central spindle, Sox21b, CG17359* (all P5) and *CG10147* (C0/P1)).

### Identifying developmental candidate genes

We next sought to test if the positional candidate genes that are expressed in the genitalia according to our RNA-seq data have a role in the development of either the posterior lobes or the claspers. To do this, we used RNAi in *D. melanogaster* to knockdown candidate genes in the smallest posterior lobe (P2) and clasper (C1) candidate regions, as well as a selection of promising genes from the other regions based on their expression profiles (Table 1, Supplementary Files 5A and 7). RNAi knockdown of the two expressed genes within P2 had no significant effect on posterior lobe size (nor on clasper size) (Supplementary File 5).

In combination with our previous study (Tanaka et al., 2015), we have now carried out RNAi for all expressed C1 candidate genes with available UAS lines (32 out of 35, Supplementary File 5). We previously observed that RNAi knockdown of *Cuticular protein 66D* (*Cpr66D*) and *Minichromosome maintenance 7 (Mcm7*) results in smaller and larger claspers, respectively (Tanaka et al., 2015). In addition to these two genes, we have now found that knocking down *hairy* (*h*), *Grunge* (*Gug*), and *fear of intimacy* (*foi*) significantly affects clasper bristle number and clasper morphology (Fig. 2A, 3E-F’ and 3J, Supplementary File 5 and 6A). Knockdown of *h* results in larger claspers with more bristles (Fig. 2A and 3F-F’), while reducing *Gug* expression gives rise to smaller claspers with fewer bristles (Fig. 2A and 3E-E’). This implies that the H and Gug TFs play opposite roles in the regulation of clasper size. Interestingly, *Gug* also appears to positively regulate posterior lobe size; since knocking down this gene significantly reduces the size of these structures (Fig. 2B and 3I-I’, Supplementary File 6A). *foi* knockdown results in severe developmental defects, with fusion of the appendages of the male external genitalia including the claspers (Fig. 2J).

**Figure 2.**
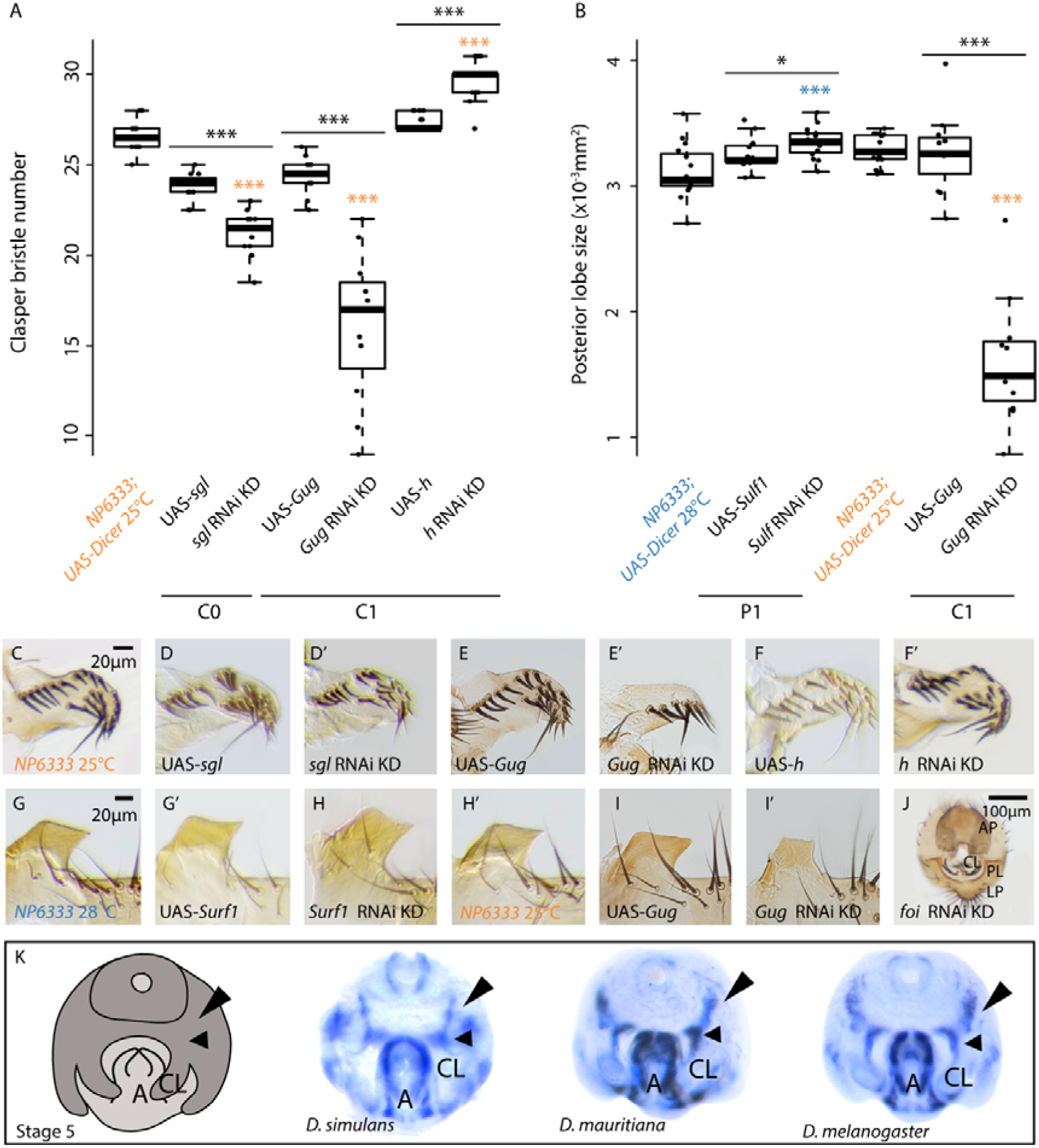
Functional analysis of positional candidate genes in *D. melanogaster* male genitalia. **(A)** Knocking down C0 candidate gene *sgl* and C1 candidate gene *Gug* resulted in significantly fewer clasper bristles compared to both the UAS (black asterisks, *p* < 0.001) and NP6333 driver controls (orange asterisks, *p* < 0.001). In contrast, knocking down C1 candidate gene *h* resulted in significantly more clasper bristles compared to the NP6333 driver (*p* < 0.001, orange asterisks) and UAS controls (*p* < 0.001, black lines and black asterisks). **(B)** P1 candidate gene *Surf1* RNAi knockdown resulted in significantly larger posterior lobes compared to both the UAS (black asterisks, *p* < 0.05) and NP6333 driver controls (blue asterisks, *p* < 0.001) (Supplementary File S6A). In addition, knocking down C1 candidate gene *Gug* resulted in the development of significantly smaller posterior lobes compared to the UAS (orange asterisks) and NP6333 driver controls (*p* < 0.001, black asterisks). Asterisks indicate significant differences detected during Tukey’s test pairwise comparisons, where *p* < 0.001 *** and *p* > 0.05 = “ns” (Supplementary File S6A). Colours indicate comparisons between the NP6333 driver control and UAS controls / gene knockdowns, while comparisons between UAS controls and knockdowns are indicated by black lines and black asterisks. Boxes indicate the range, upper and lower quartiles and median for each sample. KD = knockdown. **(C – J)** Morphology of claspers (upper row) and posterior lobes (bottom row) in NP6333 driver controls (first column), UAS controls and gene knockdowns **(D-F’** and **G’-J)**. **(K)** An illustration of stage 5 male genitalia (excluding the posterior lobes) and *in situ* hybridisations of *Cpr66D* in *Dsim w*^501^, *Dmau D1* and *Dmel w*^1118^. *Cpr66D* transcripts were detected in a wider domain along the clasper inner edge (small arrowheads) and in bands extending towards the anal plates (large arrowheads) in the two species with larger clasper. *Crpr66D* is also expressed in the aedeagus of all three species. CL = clasper primordia, A = aedeagus (internal genitalia).

**Figure 3.**
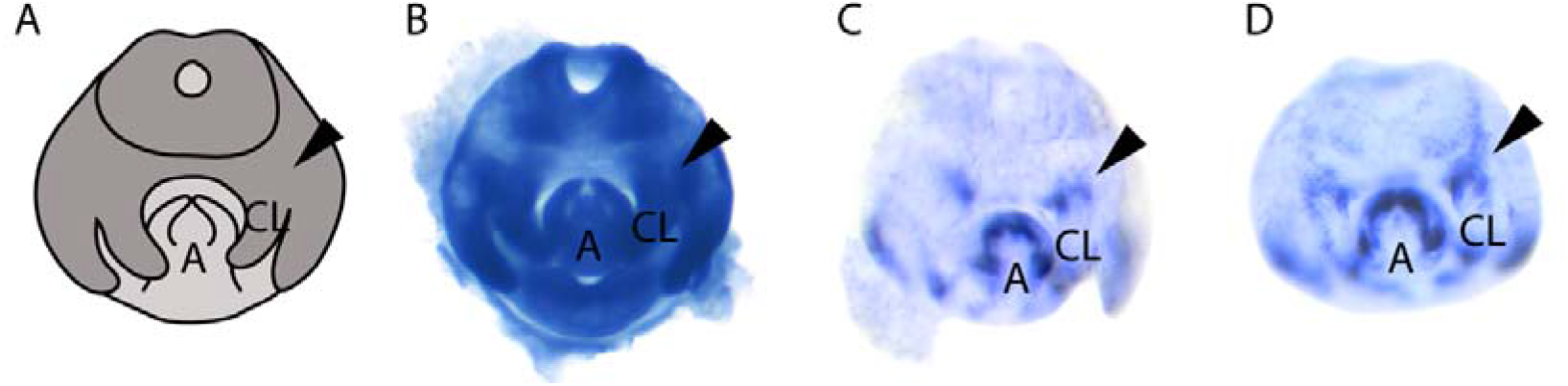
*trn* expression expands in the clasper primordia upon *h* RNAi knockdown in *D. melanogaster*. (A) A schematic of morphological development based on DAPI at Stage 5 (Hagen et al., 2019). (B) *h* mRNA *in situ* hybridisation in *D. melanogaster w^1118^*. (C) *trn* mRNA *in situ* hybridisation on UAS parental control and (D) *h* RNAi knockdown at stage 5. *h* RNAi results in a distortion of *trn* expression at the base of the claspers compared to the UAS control (black arrows). CL = claspers, A = adeagus. N = 5 for each experiment.

Therefore, the C1 region contains five promising clasper developmental candidate genes. However, only *Cpr66D* is differentially expressed between *Dsim w*^501^ and *Dmau D1* (upregulated in the latter, Supplementary File 5). Analysis of the spatial expression of *Cpr66D* during genital development revealed that this gene is expressed in a wider domain along the inner clasper edge in *D. melanogaster* and *D. mauritiana* compared to *D. simulans*, and in bands extending from this region towards the lateral edge of the anal plates (Fig. 2K).

Region C0/P1 encompasses 99 genes (Table 1). Sixty-nine of these genes are expressed in the developing genitalia according to our RNA-seq data, with twelve exhibiting significantly differential expression between *Dmau D1* and *Dsim w*^501^ (Table 1). We carried out RNAi against three of these differentially expressed genes (*Surfeit 1* (*Surf1*), *SP1173* and *CG9953*), and five other non-differentially expressed genes (*sugarless* (*and five other non-differentially sgl*), *CG32388, ventral veins lacking* (*vvl*), *CG10064* and *Lactate dehydrogenase* (*ImpL3*)). Only RNAi against *Surf1* and *sgl* affected posterior lobe and clasper development, respectively (Supplementary File 5 and 6A). RNAi against *Surf1* resulted in slightly larger posterior lobes (Fig. 2B and 2H), but had no effect on clasper bristle number (Supplementary File 5 and 6A). Interestingly, this is consistent with significantly lower expression of this gene in *Dsim w*^501^, which has larger lobes compared to *Dmau D1* (Fig. 1A, Supplementary File 5). *sgl* appears to have a role in clasper development because RNAi knockdown of this gene led to significantly smaller claspers, but had no effect on the posterior lobes (Fig. 2A, 2D-D’, Supplementary File 5).

### Interactions between genes underlying clasper divergence

*trn* is the only gene identified so far that has been shown to contribute to clasper differences between *D. simulans* and *D. mauritiana*. The *D. mauritiana* allele of *trn* generates larger claspers with more bristles than the *D. simulans* allele (Hagen et al., 2019). This is likely achieved through the expanded and/or more enduring expression of *trn* in the base of the developing claspers in *D. mauritiana* compared to *D. simulans* (Hagen et al., 2019). It was previously shown that the transcriptional co-repressor *h*, represses *trn* expression during embryogenesis in *D. melanogaster* (Chang et al., 1993, Kok et al., 2015) and in *Drosophila* Kc cells (Bianchi-Frias et al., 2004). Since we found that *h* RNAi in *D. melanogaster* results in significantly larger claspers with more bristles (Fig. 2), we hypothesised that this gene might negatively regulate clasper size through repression of *trn*.

Consistent with a previous study (Vincent et al., 2019), we found that *h* is ubiquitously expressed throughout the male genitalia of *D. melanogaster*, including the clasper primordia (Fig. 3B). We then analysed the expression of *trn* in the developing genitalia of *h* RNAi knockdowns (Fig. 3C and 3D). Upon *h* RNAi knockdown *trn* expression at the base of the developing claspers at 46 hAPF (stage 5) appears to be expanded and the edges of the domain are less well defined compared to controls, with the bands of expression extending in wisps dorsally (Fig. 3C and 3D). This ectopic expression indicates that the larger claspers produced upon *h* RNAi knockdown in *D. melanogaster* are likely due to increased *trn* expression at the base of the claspers, and that *h* acts upstream of *trn* in the clasper GRN. Despite being ubiquitously expressed throughout the male genitalia (Fig. 3B), the selective targeting of *trn* by H may explain the role of this TF in regulating clasper but not posterior lobe development; since *trn* is not expressed in the developing posterior lobes (Fig. 3C). However, *h* is expressed throughout the genital arch, and so its role in the development of the other genital structures is still unclear.

## Discussion

### Regions on chromosome 3L contributing to inter- and intra-specific variation in posterior lobe and clasper size

In this study we generated new introgressions to refine the genetic map of clasper size, clasper bristle number, and posterior lobe size differences between *D. mauritiana* and *D. simulans* on chromosome arm 3L (Fig. 1). This resulted in higher mapping resolution and indicated that some of the previously identified regions must contain multiple linked loci affecting these traits (e.g. the separate effect of regions C0 and C1 can now be distinguished compared to Tanaka et al., (2015)). As found previously, all regions identified through our introgression approach affect the claspers and/or posterior lobes consistently in the direction of their differences between the two species: *Dmau D1* DNA resulted in larger claspers and smaller posterior lobes than *Dsim w*^501^ and vice versa (Hagen et al., 2019, Tanaka et al., 2015, Zeng et al., 2000). In addition, clasper area and clasper bristle number still map to the same genomic locations, which suggests the same genes may influence both traits (Hagen et al., 2019, Tanaka et al., 2015). This could at least in part be explained by the process of bristle formation through lateral inhibition (Heitzler and Simpson, 1991) and consequently large claspers developing more bristles than small claspers. It is not clear, therefore, whether selection drove changes in clasper bristle number, and clasper size changed as a by-product, or vice versa. However, the interdigitization of clasper bristles with those of the female oviscapt would perhaps argue for the former scenario (Mattei et al., 2015).

Apart from C0/P1, all regions identified only affected either the claspers or the posterior lobes, which suggests different genes underlie the diversification in size of these two structures between *D. simulans* and *D. mauritiana* (Fig. 1). The effects observed for C0/P1 could be explained by a single evolved locus that is able to affect growth of the claspers and posterior lobes in opposite directions (with *D. mauritiana* C0/P1 alleles generating smaller posterior lobes and larger claspers (see Fig. 1B and 1C)). However, since C0/P1 is still a relatively large region, it is likely that further mapping would resolve this region into distinct clasper and posterior lobe loci. Genes within region C0/P1 may underlie intra-specific variation as well as interspecific differences in posterior lobe size. This region overlaps with the 3L QTL peak observed in other inter-specific mapping studies of differences in posterior lobe size between *D. simulans* and *D. mauritiana* or *D. sechellia* (Liu et al., 1996, Macdonald and Goldstein, 1999, Masly et al., 2011, Zeng et al., 2000), as well as QTL peaks found in studies that mapped genetic variation underlying differences in posterior lobe size between *D. melanogaster* strains (Hackett et al., 2016, McNeil et al., 2011, Takahara and Takahashi, 2015). Therefore, P1 posterior lobe candidate genes, such as *Surf1*, represent excellent candidates for contributing to variation in the size of this structure within and between species.

### Genome-wide gene expression during genital development in D. mauritiana and D. simulans

We carried out RNA-seq to identify and compare genes expressed in the developing genitalia between *D. mauritiana* and *D. simulans*. As well as allowing us to filter positional candidates, this also provided a genome-wide perspective of gene activity during genital development as well as differential expression between species.

We detected a total of 8984 genes in the developing genital arch of *Dsim w*^501^, and 8458 genes in that of *Dmau D1* (Supplementary File 6A). This included all the key genes known to pattern the genital disc, such as homeotic genes and sex-determination genes (Casares, 1997, Chen and Baker, 1997, Estrada et al., 2003, Estrada and Sanchez-Herrero, 2001, Keisman and Baker, 2001, Sanchez and Guerrero, 2001) and signalling genes, including *wingless, decapentaplegic* and *hedgehog* (Abdelilah-Seyfried et al., 2000, Casares, 1997, Chen and Baker, 1997, Keisman and Baker, 2001, Sanchez and Guerrero, 2001), as well as the TFs as *cubitus interruptus, engrailed* (Eaton and Kornberg, 1990, Kornberg et al., 1985, Simmons and Garcia-Gonzalez, 2011), *dachshund* (Keisman and Baker, 2001), *Distal-less* (Estrada and Sanchez-Herrero, 2001) and *Drop* (Chatterjee et al., 2011).

We found 676 and 143 genes that are exclusively expressed in the developing genitalia of either *Dsim w*^501^ or *Dmau D1*. The *Dsim w*^501^ male genital specific genes are enriched for iron ion binding proteins, while *Dmau D1 are* enriched for multiple IRs. IRs are a conserved family of chemosensory receptors known for their role in olfaction (Benton et al., 2009, Grosjean et al., 2011, Min et al., 2013, Silbering et al., 2011, Ziegler et al., 2013). Interestingly, some IRs, for example, *IR52c* and *IR52d*, are candidate taste and pheromone receptors (Koh et al., 2014) that are expressed in a sexually dimorphic manner on the sensilla of the *D. melanogaster* male foreleg, which makes contact with the female during courtship (Koh et al., 2014). The neurons in which these IRs are expressed in *D. melanogaster* males are only activated upon contact with females of the same species (Koh et al., 2014). Therefore, the differences in IR expression between the genitalia of males of different *Drosophila* species may be an evolved mechanism to prevent conspecific mating.

Of the genes that are expressed in both *Dsim w*^501^ and *Dmau D1*, we found that 1169 were differentially expressed (Supplementary File 6). This includes 61 out of the 802 TF encoding genes, five of which have been shown to be expressed in both the claspers and the posterior lobes of *D. melanogaster* (*hinge3, Myb oncogene-like, single stranded-binding protein c31A, Sox21b* and *Enhancer of split m3, helix-loop-helix*) (Vincent et al., 2019). This suggests that the regulatory landscape of developing genitalia is generally conserved between *D. mauritiana* and *D. simulans*. However, the differentially expressed TFs will help to better understand the gene regulatory networks involved in genital development and evolution, and represent excellent candidates genes for further investigation.

### Functional analysis of expressed positional candidates on chromosome 3L during genital development

Our mapped regions on chromosome 3L encompass 260 genes that are expressed during male genital development in *Dsim w*^501^ and *Dmau D1*, of which 50 are differentially expressed (Table 1). We have now analysed the function of 58 of the expressed genes by RNAi knockdown in *D. melanogaster*, including 32 out of the 35 genes expressed in C1 (including those we studied previously in Tanaka et al. (2015)), as well as all expressed genes in P2 (Table 1). Note that we did not just focus on differentially expressed genes because genes can exhibit localised differences in expression during genital development that may contribute to morphological differences (Hagen et al., 2019).

RNAi against the expressed P2 genes did not have any significant effect on the posterior lobes (Supplementary File 5A). While RNAi against these genes simply may not have worked, alternatively a non-protein coding element in this region could underlie the phenotypic effect of region P2 on posterior lobe size. P2 contains a microRNA, *mir-4940*, as well as the long noncoding RNA CR45408 (Thurmond et al., 2018). Therefore, the causative element in P2 could be either of these factors or even a long-range enhancer responsible for the differential regulation of a gene outside P2 between these two species.

Our functional analysis of region C0/P1 identified two excellent candidate genes *Surf1* and *sgl*. *Surf1*, which encodes a nuclear gene, appears to negatively regulate posterior lobe size and is expressed more highly in *D. mauritiana* than in *D. simulans*, consistent with the RNAi resulting in larger posterior lobes (Fig. 2A and 2H). *sgl* has been implicated in boundary formation and may interact with Wnt signalling (Hacker et al., 1997). RNAi against *sgl* resulted in smaller claspers (Fig. 2A and 2D’), but this gene is not differentially expressed between *D. mauritiana* and *D. simulans*. However, since C0/P1 is a large region that is likely to contain many other developmental candidates, higher resolution mapping and functional analysis of genes in C0/P1 is needed.

We have now studied the function of 32/35 candidate genes in C1 during genital development (Tanaka et al., 2015). This revealed five interesting candidate genes for clasper development and evolution: *Gug*, *foi*, *Mcm7*, *Cpr66D* and *h*. However, only *Cpr66D* is differentially expressed between *Dsim w*^501^ and *Dmau D1*, and expression of this gene is more extensive along the inner edge of the claspers and in bands extending towards the anal plates in *Dmau D1* compared to *Dsim w*^501^ (Fig. 2K). *Cpr66D* encodes a structural protein that forms chitin-based cuticle (Chandran et al., 2014, Ren et al., 2005, Stahl et al., 2017) and its role in genital development merits further study.

We also found evidence for potential regulatory interactions between genes in mapped regions during genital development. Repression of *trn* by H has been predicted (Bianchi-Frias et al., 2004, Kok et al., 2015) or shown (Chang et al., 1993) in different developmental contexts. We found that H also negatively regulate *trn* expression in the developing claspers of *D. melanogaster;* with the larger claspers resulting from *h* RNAi likely being caused by the consequential expansion of *trn* expression (Fig. 2A, 2F’ and 3D). H also negatively regulates *trn* expression during embryogenesis to help define compartmental boundaries (Chang et al., 1993; Pare et al., 2019). Therefore this regulatory interaction could represent a more general mechanism for co-ordinating the correct positioning of cells during development. However, *h* is not differentially expressed between *Dsim w*^501^ and *Dmau D1* and appears to be ubiquitously expressed in the developing genitalia of *D. melanogaster* (Fig. 3B) (Vincent et al., 2019). Although it is possible that there could also be localised differences in *h* expression in the developing genitalia, these observations suggest that the differences in *trn* expression between *Dsim w*^501^ and *Dmau D1* could be the result of protein-coding changes that affect the DNA-binding efficiency of H, or variation in the number and/or sensitivity of H binding sites in *trn* regulatory elements. Indeed, there are several predicted H binding sites across the *trn* locus (data not shown), but identification of *trn* genital enhancers and further analyses of H binding sites between *D. mauritiana* and *D. simulans* is needed to test this further.

In addition to *trn*, H may regulate multiple genes during clasper development including candidates revealed by our mapping and functional analyses. For example, H is also predicted to negatively regulate the C1 candidate gene, *Gug* (Yeung et al., 2017). Indeed, Gug itself is predicted to regulate the C0 candidate gene *sgl*, as this gene contains a Gug binding site in its intron (Yeung et al., 2017). However, since Gug acts as a transcriptional co-repressor, and RNAi against both *Gug* and *sgl* reduces clasper size, it is unclear at this stage if this is there is a regulatory interaction between these genes in the developing claspers. It will be interesting to test these predictions in the future to learn more about the architecture of the gene regulatory network for clasper development and how this evolved during the rapid diversification of these structures.

## Materials and Methods

### Introgression line phenotyping

We generated new recombinants in our candidate regions by backcrossing virgin *IL D11.01* / *Dsim w*^501^ heterozygous females, and virgin *IL D08.04* / *Dsim w*^501^ heterozygous females to *Dsim w*^501^ males. *IL D1101* is an introgression line with *D. mauritiana D1* DNA in the genomic location 3L: 7527144..15084689 Mb and encompasses the candidate regions C1, P2 and P3 (Tanaka et al., 2015). *IL D08.04* is an introgression line with *D. mauritiana w*-DNA on 3L: 5911371..9167745 Mb (R2.02 *D. simulans*) and includes candidate regions P1 and C1 (Tanaka et al., 2015). New recombinants were detected by selecting for the loss of the visible marker D1 (Tanaka et al., 2015, True et al., 1997) (Fig. 1D), restriction fragment length polymorphisms and sequencing markers (see Supplementary File 7 for primer list). New introgression lines (Supplementary File 1) were all maintained as homozygous stocks.

Male genitalia were phenotyped from flies cultured under controlled growth conditions. All males used were progeny of 10 females and five males that were transferred every two days, and allowed to develop at 25°C in a 12 hours light/dark cycle incubator unless otherwise stated. All adult males were then kept on a standard cornmeal diet at 25 °C for at least three days before collection and storage in 70% EtOH.

Where possible, two or three replicates of ILs were phenotyped. Replicates are defined as introgression lines derived from the same recombination event and therefore containing the same introgressed region of *D. mauritiana* DNA. The abdominal tip and T1 leg were dissected for each fly in 70% EtOH, and transferred to Hoyer’s medium. Using entomological pins, the posterior lobes were then dissected away from the claspers and anal plates. The claspers, posterior lobes and T1 tibia were mounted in Hoyer’s medium for imaging.

Images were taken using a Zeiss Axioplan light microscope at 250X magnification for the claspers and lobes and 160X for the T1 tibia, using a DFC300 camera. Clasper area, posterior lobe size and tibia length were measured manually using ImageJ (Schneider et al., 2012), and bristle number was counted for each clasper (Supplementary File 2A). T1 tibia length was used as a proxy for body size, in order to assess consistency in rearing conditions and to ensure genital differences were not a result of general differences in size. Most introgression lines showed no significant difference in T1 tibia length compared to *Dsim w*^501^ (Supplementary File 2G), and since genitalia are hypoallometric (Coyne et al., 1991, Eberhard, 2009, Liu et al., 1996, Macdonald and Goldstein, 1999, Masly et al., 2011, Shingleton et al., 2009), the phenotypic data was not corrected for body size. A detailed description of statistical methods and the comparisons used to map candidate regions based on these data can be found in the Supportive Text.

### RNA sequencing and differential expression analysiss

Three independent RNA-seq library replicates were generated for *Dsim w*^501^ and *Dmau D1* developing male genitalia. Flies were reared under the above conditions, and white pre-pupae collected. Males were selected using gonad size and allowed to develop in a humid container at 25°C until either stage 2 or stage 4.5 for extraction (see staging guide in Hagen et al., (2019)). Between these stages, the claspers develop from a ridge structure to a distinct appendage separate from the surrounding tissue, and the posterior lobe has begun to extend outwards from the lateral plate primordia (Hagen et al., 2019). The heads of pupae were impaled with a needle onto a charcoal agar plate and submerged in 1xPBS. Dissection scissors were used to remove the distal tip of the pupal case and the outer membrane, and pressure applied to the abdomen to allow the developing genitalia to be quickly expelled from the pupal case and dissected away from the abdomen. Note that the entire genital arch, including internal genital organs (but not including abdominal tissue), was isolated for RNA extraction. The genitalia from fifteen males from the two stages were collected and then combined in TRIzol. RNA was then extracted using standard procedures. Quality and quantity of RNA was verified using a Qubit. Samples were sequenced by the NERC Biomolecular Analysis Facility (NBAF) at the Centre for Genomic Research, University of Liverpool, where dual-indexed, strand-specific RNA-seq libraries were prepared using NEBNext polyA selection and Ultra Directional RNA preparation kits. Samples were then sequenced using Illumina HiSeq 4000 (paired-end, 2×150 bp sequencing). *Dsim w^501^ and Dmau D1* reads were mapped against reannotated reference coding sequences (Torres-Oliva et al., 2016). Raw fastq files are available upon request. Genes were considered to be expressed if transcripts per million (TPM) > 1 in all three biological replicates. Genes were only considered differentially expressed in comparisons where padj (FDR) < 0.05.

### Gene ontology analysis

In order to investigate the nature of the expressed, not expressed and differentially expressed genes in our RNA-seq dataset, we determined their ontology using PANTHER (Thomas et al., 2003). We conducted Overrepresentation Tests (Released 20190711) of gene ontology (released 09/12/2019) for the positional genes against the *D. melanogaster* reference list using the Fisher Test (Thomas et al., 2006). Genes were considered significantly overrepresented when padj (FDR) < 0.05.

### Annotation of transcription factors present in RNA-seq data

In order to extract the genes encoding TFs from the RNA-seq dataset, we used the databases of genes from Flymine (https://www.flymine.org/flymine/begin.do (Lyne et al., 2007), amiGO (http://amigo.geneontology.org/ (Carbon et al., 2009)) and Flybase (Thurmond et al., 2018) and bioinformatic analysis and manual curation from Hens et al. (2011). We filtered the genes in our dataset corresponding to TFs by their GO terms and gene groups in molecular function using the previously mentioned sources. The GO terms used were the following: ‘FlyTF_putativeTFs’ from Flymine (Lyne et al., 2007), ‘Transcription factor regulator activity’ and ‘Transcription factor coregulator activity’ from amiGO (Carbon et al., 2009), ‘Transcription factor gene group’ and ‘transcription coregulator activity’ from Flybase (Thurmond et al., 2018) and the dataset of TFs from Hens et al., (2011). Genes that were annotated with these terms in any of the four resources were considered TF or co-factor encoding genes and used for downstream analysis.

### RNAi knockdown of candidate genes

The developmental role of genes was tested using RNAi in *D. melanogaster*. UAS-RNAi lines for these genes were ordered from both Vienna *Drosophila* RNAi centre and TRiP lines from Bloomington stock centre (see Supplementary File 6A for stock numbers). UAS males of our candidate genes were crossed to *NP6333–Gal4* (“*NP6333*”) driver virgins (*P[GawB]PenNP6333*) (Chatterjee et al., 2011) carrying *UAS-Dicer-2 P[UAS-Dcr-2.D]*. RNAi knockdown was conducted at either 25°C or 28°C (Supplementary File 6A), under identical rearing conditions, and dissection, imaging and analysis was carried out as described above (Supplementary File 6B). To assess the role of a gene during genitalia development, we compared the phenotype of genital structures of gene knockdowns against the respective *NP6333* driver controls using a Dunnett’s test (Supplementary File 6A). If the gene knockdown phenotype differed significantly from the NP6333 driver control, we then assessed whether or not this significant effect is a result of genetic background (e.g. an effect of the UAS-parental phenotype), or reflects a role of the gene in genital development. To do this, we compared all three experimental groups of males using an ANOVA (Supplementary File 6A). If this was significant, we then analysed where these differences arise from using a Tukey’s test, and only concluded genes have a developmental role in the genitalia if the RNAi knockdown males were significantly different in phenotype compared to both parental controls.

### In situ hybridisation

Sample collection, RNA extraction, cDNA synthesis and probe synthesis were conducted as described in Hagen et al., (2019). We performed in situ hybridisation to detect expression of *Cpr66D* in *D. mauritiana, D. simulans* and *D. melanogaster, h* in *Dmel W*^1118^ and *trn* in *UAS-h* Bloomington TRiP 27738, *NP6333-Gal4;UAS-Dicer* x *UAS-h* Bloomington TRiP 27738 using species-specific probes. Probes were generated using the following oligos (forward followed by reverse) with the addition of T7 linker sequences added to the 5’ end of each primer; *trn* (514 bp) ATCGAGGAGCTGAATCTGGG and TCCAGGTTACCATTGTCGCT (Hagen et al., 2019), *Cpr66D* (314 bp) CTCCTCGTATCAGTTTGGCTTC and CTGGTGGTACTGTGGCTGCT. Antisense *h* probes were generated by amplifying from a BLUESCRIBE plasmid that contained sequences for all three *h* coding exons using T7 primers (a gift from B. Jennings, Oxford Brookes University). In situ hybridizations were based on the Carroll lab *“Drosophila* abdominal *in situ”* protocol (http://carroll.molbio.wisc.edu/methods.html) with minor modifications.

## Supporting information

Supportive text

## Author contributions

JFDH, CCM, APM, and MDSN designed this project; JFDH, CCM, JFJ, FAF, LBG, SRB, KMT, AMR, and SA performed the experimental work; JFDH, CCM, APM, KMT., and MDSN analysed data; and JFDH, APM., and MDSN wrote the paper.

## Acknowledgements

This work was funded by a NERC grant (NE/M001040/1) to APM, and a JSPS KAKENHI (15J05233) grant to KMT. JFDH and JFJ were funded by Nigel Groome studentships from Oxford Brookes University, and SRB and AMR by BBSRC DTP studentships. We thank the NERC Biomolecular Analysis Facility (NBAF) at the Centre for Genomic Research, University of Liverpool for sequencing.

