## Supplementary material for "Unravelling the genetic basis for the rapid diversification of male genitalia between *Drosophila* species": Supportive text

**Statistical analysis and effect size calculations for introgression mapping**

To increase the resolution of clasper and posterior lobes candidate regions, we generated 23 new introgression lines (ILs) in the genomic region 3L: 5911371-15084689 Mb (Supplementary File 1). Phenotypic measurements and descriptive statistics for each introgression line are shown in Supplementary File 2A and 2B respectively. First, we tested for normality of the data generated from each introgression line using Shapiro-Wilk test (Supplementary File 2C). Most phenotypes were normally distributed data for each line, except for clasper bristle number, which was often non-normal in distribution (Supplementary File 2C). Since many introgressed regions were represented by multiple lines, we first tested for significant differences between introgression line replicates. If replicates were non-normally distributed (mostly clasper bristle number measurements), we conducted a Kruskal-Wallis test, followed by a Pairwise Wilcox test. Replicate lines that contained normally distributed data were compared using an ANOVA followed by a Tukey’s test. For introgressed regions represented by two replicates that were non-normally distributed, we conducted a Pairwise Wilcox test, or for those that were normally distributed phenotypes were compared using a Welch Two-Sample t-test. If the replicate lines were not significantly different from each other, they were considered equivalent and the phenotypic measurements were pooled (Supplementary File 2C), replicates that could not be merged are highlighted in blue for each trait, and were omitted from downstream mapping analysis. To assess the effect of the introgressed region, these pooled values were first compared against the parental *Dsim w*^501^ line. For traits which yielded normally distributed data (clasper size, posterior lobe, tibia measurements), this was achieved using an ANOVA followed by a Dunnett’s test (Supplementary File S2D - 2D). For those that were not normally distributed (clasper bristle number), a Kruskal-Wallis Dunn’s test was performed. If the genital phenotype of an introgression line was significantly different from that of *Dsim w*^501^, we concluded that there must be at least one locus in the introgressed region affecting the phenotype of the trait. The phenotype of overlapping introgression lines was then compared; for lines containing normally distributed phenotypic data, we conducted an ANOVA, while data that were not normally distributed were compared using a Kruskal-Wallis test. If there were no significant differences between lines overlapping, then the minimal shared region of overlap was used to define the newly resolved candidate region. If significant differences were found between overlapping lines, we concluded the existence of multiple causative loci. In order to determine the location of these loci, we then conducted pairwise comparisons between the lines. For normally distributed data, lines were compared using a Tukey’s test, while those that were not normally distributed data were compared using a Pairwise Wilcox test (Supplementary File S2D – 2G). All statistics were conducted in version 1.2.1335 RStudio (RStudioTeam, 2018).

After determining the location of the resolved candidate regions, the phenotypic effect size of each region was calculated. This was defined as the proportion of phenotypic difference in clasper size between *Dsim w*^501^ and *Dmau D1* that the evolved loci can explain. This value was calculated as a percentage, based on the average phenotype of lines containing introgressed *D. mauritiana* DNA including the candidate region concerned.

**Posterior lobe size mapping**

Based on the ILs with no significant effect on posterior lobe size compared to *Dsim w*^501^ (*IL D21.43, ILD21.43e and IL 8.15*), we excluded 3L: 7441137..8903265 and 3L:14402074..15084689 from our posterior lobe map.

*Defining the P1 region*

The new *IL 6* has posterior lobes that are significantly smaller than those of *Dsim w*^501^ (Dunnett’s test, p < 0.001, Supplementary File 2F). This line contains introgressed DNA from *D. mauritiana* between 3L: 5911371..6852222, and does not differ significantly in posterior lobe size from other overlapping ILs (Fig. 1B and Supplementary File 3C) (Tukey’s test, p > 0.05) except for *IL D08.04*, which has even smaller posterior lobes (Tukey’s test, *p* < 0.001). The introgressed region in *IL D08.04* (3L: 5911371..9167745) partially overlaps with that of *IL 6* (Supplementary File 1). These results suggest that *IL 6* and *IL D08.04* contain posterior lobe candidate region P1 in their region of overlap, but that *IL D08.04* contains an additional region, P2. P1 is therefore defined maximally by the breakpoints of *IL 6* (3L: 5911371..6852222). Based on the phenotypic effect of lines that contain no other posterior lobe candidate regions (*IL 6, IL 137, IL 21, IL 7*), we estimate that P1 contributes to 3.7% of posterior lobe size differences between *Dsim w*^501^ and *Dmau D1* (Table 1).

*Defining the P2 region*

P2 must be located in the region between the right-hand breakpoints of *IL D08.04* and *IL 6*. Since *IL 7* breaks in this region and has posterior lobes that are even larger than *IL D08.04* (Tukey’s test, p < 0.001, Supplementary File 3C), P2 must lie between the right-hand breakpoints of *IL 7* and *D08.04* (3L: 8096301..9167745). However, as previously mentioned, part of this region can be excluded (3L: 7441137..8903265) because other introgressions with DNA in this region (*IL D21.43e* and *IL D21.43*) are not significantly different in posterior lobe size compared to *Dsim w*^501^ (Supplementary File 2F). The remaining candidate P2 region, between 3L: 8903265..9167745, overlaps with the left-hand breakpoints of *IL D11.01, IL 4.6* and *IL 4.11* (Supplementary File 1)*. S*ince the posterior lobes of *IL D11.01* males are smaller than those of *IL 4.6* and *IL 4.11* males, the left-hand break points of the latter introgressions must define the end of P2 (Supplementary File 3C). The common region highlighted by these comparisons, 3L: 8903265..9158500 (i.e. between the right-most breakpoint of *IL D21.43* and the left-most break point of ILs *4.11* and *4.6*), therefore contains the newly resolved P2. Based on the average effect size differences between *IL D11.01, IL 4.6* and *IL 4.11, and IL 7* and *D08.04 ,* we predict the P2 region explains 6% of differences in posterior lobe size between the species (Table 1).

*Defining the P3 region*

*IL 11.10* and *IL 4.12* have the same right-most breakpoint of introgressed *Dmau D1* DNA, and only differ in their left-most breakpoint (3L: 9178292..9572682, Supplementary File S1). Since *IL 11.10* males have significantly larger posterior lobes than both *IL 4.6* and *IL 4.11* (Tukey’s test, p < 0.001), but *IL 4.12* males have statistically the same sized posterior lobes as *IL 4.6* and *IL 4.11* (Supplementary File 3C), we predict posterior lobe candidate region P3 lies within the left-most break point of these lines. Based on the effect size differences between *IL 4.11, IL 4.6, IL 11.10 and IL 4.12*, we predict P3 contains variation that explains 5.9% of posterior lobe size differences between the species (Table 1).

*Defining the P4 region*

The introgressed *D. mau D1* DNA in *IL 16.30* significantly reduced posterior lobe size compared to *Dsim* w*^501^* (Dunnett’s test, *p* < 0.001, Supplementary File 2F). Since we have no other IL that contain introgressed DNA that breaks in the region in order to resolve it (Supplementary File 1), we mapped region P4 to the boundaries of *IL 16.30* (i.e. 3L: 12277961..12737959). Based on the effect size of IL’s containing this introgressed region (*IL 10, IL 4.14, IL 43, IL 16.14 and IL 16.30*), we predict that P4 contains variation explaining 4.9% of posterior lobe size differences between the species (Table 1).

*Defining the P5 region*

Since *IL 11.10, IL 164* and *IL 111* have significantly smaller lobes than both *IL 10.15* and *IL 8.15* (Tukey’s test, *p* < 0.001, Supplementary File 3C), P5 maps within the *D mau D1* DNA shared by *IL 11.10, IL 164, IL 111,* but not shared by *IL 10.15* and *IL 8.15* (Fig. 1D). Therefore, P5 lies in 3L: 13393862..14532063, and based on the differences in effect size between *IL 11.10, IL 164, IL 111, IL 10.15* and *IL 8.15,* contributes to 9% of posterior lobe size differences between the species (Table 1).

**Clasper bristle number and clasper size mapping**

Five of the introgression lines we analysed had no significant effect on clasper size or bristle number compared to *Dsim w*^501^ (*IL 8.6, IL 8.15, IL 16.30, IL 82, IL 16.12*, Supplementary File S2D and 2E). Therefore, regions 3L: 12277961..12911597 and 3L: 14402074..15084689 were excluded from our clasper mapping.

*Defining the C0 region*

*IL 6* contains introgressed *Dmau D1* DNA that contributes to significantly larger claspers compared to *Dsim w*^501^ (Dunnett’s test, *p* < 0.001, Supplementary File S2C), with more bristles (Dunn’s test, *p* < 0.01, Supplementary File S2D) (Fig. 1C). Therefore, clasper candidate region C0 maps to the same genomic region as P1, 3L: 5911371..6852222. Based on the phenotypic effect of lines that contain no other clasper candidate regions (*IL 6, IL 137, IL 21, IL 7, IL 56*), we predict C0 explains 10.8% of clasper size differences, and 24.8% of clasper bristle number differences between *D. simulans* and *D. mauritiana* (Table 1)*.*

*Defining the C1 region*

*IL D21.43e* contains larger sized claspers (Dunnett’s test, p < 0.001, Supplementary File 2D), and more bristles (Dunn’s test, p < 0.001, Supplementary File 2E) than *D. simulans w*^501^*.* Since *IL D21.43e* is not significantly different in clasper size or bristle number compared to *IL D08.04, IL 7, IL D23.42* or *IL D21.43, C1* can be refined to the region of *Dmau D1* DNA shared by all of these lines (Fig. 1D)*.* The smallest overlapping region of these lines lies between the left-most break point of *IL D23.42* and the right-most break point of *IL D21.43e*.

Therefore, none of the newly generated ILs could be used to resolve previously predicted candidate region C1 (Supplementary File 1)*,* since our analysis maps C1 to the same breakpoints as previous (Tanaka et al., 2015). Therefore, C1 has been mapped to the same genomic location as previous studies (Tanaka et al 2015); to a region between 3L: 8253548..8699816. Based on the effect sizes of *IL D08.04, IL 7, IL D23.42, IL D21.43* and *IL D21.43e*, C1 contains variation accounting for 6% of clasper size differences, and 21% of clasper bristle number differences between the species (Table 1).

Interestingly, although *IL D08.04* contains introgressed *Dmau D1* DNA encompassing both C0 and C1 (overlapping entirely with both *IL D21.43e* and *IL 6*), *IL D08.04* is not significantly different in clasper size or bristle number compared to either of these lines (Supplementary File 3A and 3B). This indicates that the loci in C0 and C1 do not act additively to affect clasper morphology.

*Defining the C2 region*

The coordinates of region C2 were reported previously and this region lies at 3L:12734279..12911597 (Hagen et al., 2019) (Fig. 1).
